# Efficient enrichment of synchronized mouse spermatocytes suitable for genome-wide analysis

**DOI:** 10.1101/2022.01.11.475957

**Authors:** Agustin Carbajal, Irma Gryniuk, Rodrigo O. de Castro, Roberto J. Pezza

## Abstract

Chromatin-based mechanisms regulating developmental transitions during meiosis are fundamental but understudied aspects of male gametogenesis. Indeed, chromatin undergoes extensive remodeling during meiosis, leading to specific patterns of gene expression and chromosome organization, which ultimately controls fundamental meiotic processes such as recombination and homologous chromosome associations. Recent game-changing advances have been made by analysis of chromatin binding sites of meiotic specific proteins genome-wide in mouse spermatocytes. However, further progress is still highly dependent on the reliable isolation of sufficient quantities of spermatocytes at specific stages of prophase I. Here, we describe a combination of methodologies adapted for rapid and reliable isolation of synchronized fixed mouse spermatocytes. We show that chromatin isolated from these cells can be used to study chromatin binding sites by ChIP-seq. High quality data we obtained from INO80 ChIP-seq in zygotene cells was used for functional analysis of chromatin binding sites.

## Introduction

Gametogenesis is the developmental program that sexually reproducing organisms utilize to produce gametes with half the somatic chromosome number. This is achieved through meiosis, in which two rounds of chromosome segregation follow a single round of DNA replication. Meiosis involves highly regulated cell developmental transitions, which are coordinated at chromatin level and ultimately control proper reproductive tissue development and sustained germ cell production in adults. For example, during the meiotic prophase I spermatocytes undergo extensive changes in the patterns of gene expression supporting a series of unique events including pairing, synapsis, and recombination of chromosomes, to ultimately ensure their accurate segregation. These processes are coordinated and controlled at the chromatin level by regulators such as chromatin remodeling complexes, which meiotic functions in any organism are still poorly understood.

A key task for researchers working in various aspects of meiosis at the genome wide level is obtaining enough high-quality spermatocytes enriched at any of the different stages of prophase I for use in assays such as ChIP-seq and RNA-seq. An important limitation in this endeavor is that meiotic cells are asynchronous - adult testis primary spermatocyte population can be found as a combination of all meiotic stages from pre meiotic S-phase to diplotene. Recent breakthrough methodology (1,2) allows synchronization of spermatocytes in testis. In this approach, spermatocyte maturation is arrested at the stage of undifferentiated spermatogonia on the newborn mouse by administration of fertilysin (WIN18,446), a compound that inhibits the synthesis of retinoic acid. Following this treatment, an injection of retinoic acid induces gametogenesis progression, resulting in the generation of synchronously maturing population of spermatocytes. The time encompassed between retinoic acid administration and cell harvest determine the stage of prophase I in which most spermatocytes are found (Fig. 1A).

**Figure 1.**
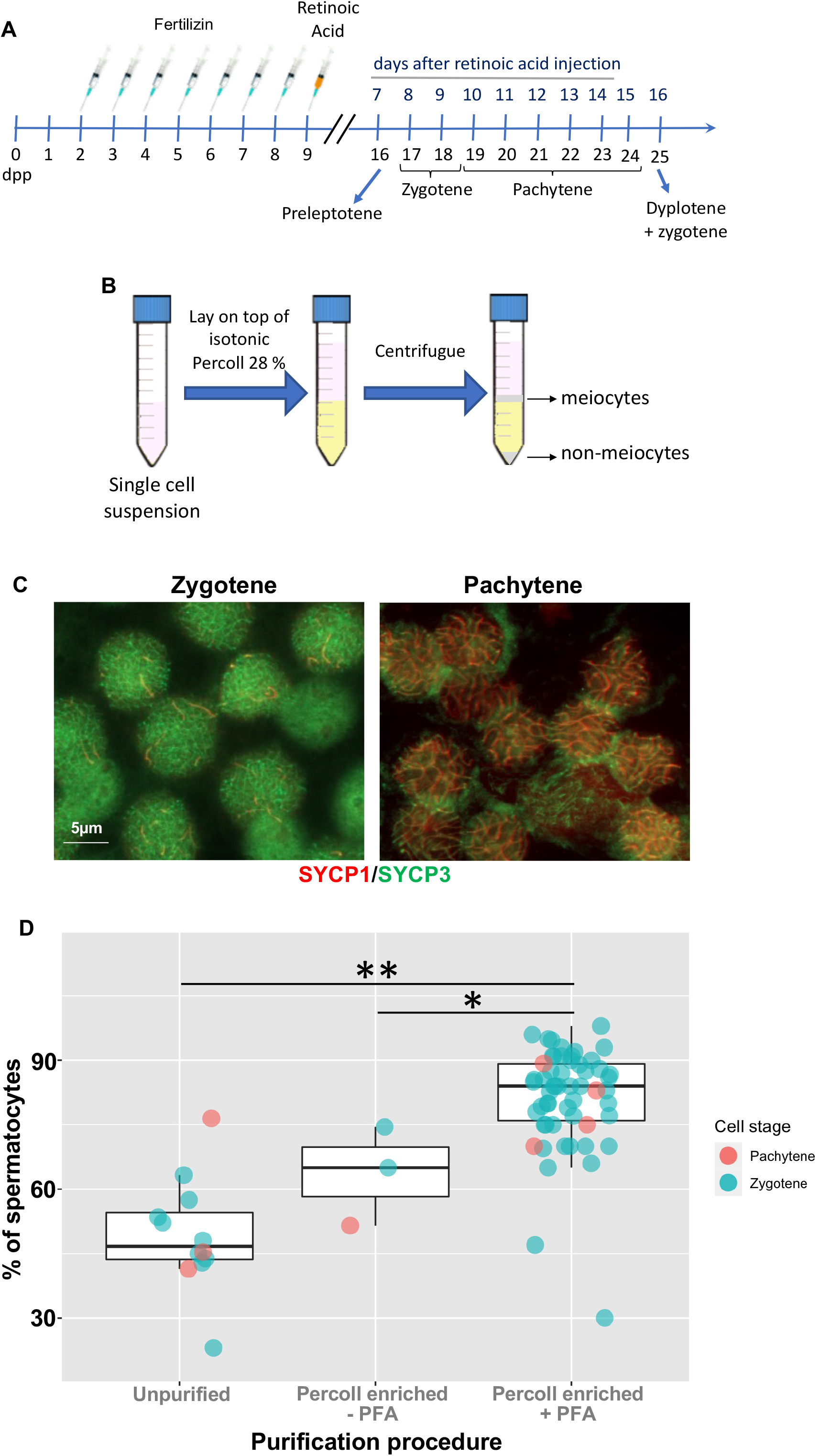
Enrichment of synchronized mouse spermatocytes. **A.** Schematic showing neonate mice treatment for synchronization of spermatocytes and collection of cells at different stages of prophase I. **B.** Diagram representing PFA-fixed spermatocyte centrifugation on a discontinuous Percoll gradient. **C.** Representative image showing zygotene and pachytene spermatocytes after Percoll gradient. **D.** Percentage of spermatocytes before Percoll gradient (mean ± standard deviation, 49.4 ± 13.1 %, n = 12), purified using a discontinuous Percoll gradient without cell fixation (“-PFA”, 63.7 ± 11.6 %, n = 3) or purified after fixing cells (“+PFA”, 81.3 ± 11.8%, n = 55). Tukey’s significance test, p < 0.05 (*) or < 0.001 (**).

An important challenge after spermatocyte synchronization is the isolation of the spermatocytes from other cell populations such as spermatogonia and somatic cells. This problem has been addressed by FAC sorting of synchronized spermatocytes (1,3,4). However, FACS requires specialized equipment and technical expertise, and is time consuming when a substantial number of cells are required for experimentation, which is often the case for genome wide analysis approaches. Here, we describe a rapid, simple, and robust approach to obtain enriched fractions of synchronized mouse spermatocytes at different stages of prophase I. In our approach we employ centrifugation of single cell suspension on a discontinuous percoll gradient, from which we obtain high yield of enriched spermatocytes in a relatively short time. Zygotene cells obtained using this method were used to perform INO80 ChIP-seq, a chromatin remodeler with important functions in meiosis. Our report suggests a possible link between this important chromatin-based regulator to critical meiotic processes.

## Materials and methods

### Mice

We used F1 hybrids mice generated by crossing DBA/2J × C57BL/6J. Mice were obtained from the Jackson Laboratory and bred in-house. All experiments conformed to relevant regulatory standards guidelines and were approved by the Oklahoma Medical Research Foundation-IACUC (Institutional Animal Care and Use Committee, protocol 20.33).

### Cytology

We employed established experimental approaches for the visualization of chromosomes in chromosome surface spreads (5). Incubations with primary antibodies (SYCP3 (6), SYCP1 (Novus, NB300-229), and γH2AX (Millipore, 05-636)) were carried out for 16 h at 4 °C in 1 X PBS plus BSA 2.5 % as we previously described (6,7). Following three washes in 1 X PBS, slides were incubated for 1 h at room temperature with secondary antibodies. A combination of fluorescein isothiocyanate (FITC)-conjugated goat anti-chicken IgG (Jackson laboratories), Tetramethylrhodamine (TRITC)-conjugated goat anti-rabbit IgG, and Cy5-conjugated anti mouse IgG, each diluted 1∶300, was used for simultaneous immunolabeling. Slides were subsequently counterstained with 2 μg/ml DAPI containing Vectashield mounting solution (Vector Laboratories) and sealed with nail varnish. We used Zen Blue (Carl Zeiss, Inc.) for imaging acquisition and processing.

### Spermatocyte synchronization

Spermatocytes at prophase I were synchronized as described in (1). Briefly, starting at 2 dpp mice were injected for seven consecutive days with fertilysin (MP Biomedicals, cat. # 158050) to arrest gametogenesis at the spermatogonia stage. The next day (9 dpp), mice were injected with retinoic acid (Sigma, cat. # R2625) to induce their coordinated maturation (Fig. 1A). To collect zygotene and pachytene cells, mice were euthanized at 8 and 10 days after retinoic acid injection, respectively.

### Enrichment of primary spermatocyte by centrifugation on discontinuous Percoll gradient

Primary spermatocytes were purified from synchronized mice testes using the following protocol (indicated amounts have been adjusted to two mice, four testes). Note that at some steps the protocol has two variants, “F” or “NF” which stand for “PFA-<u>F</u>ixed” and “<u>N</u>on-<u>F</u>ixed” respectively. **1-** After detunication, testes were placed in a 15 ml conical tube containing 2 ml of prewarmed (35 °C) collagenase solution (1 mg/ml type I collagenase (Worthington, cat. # LS004196) and 60 U/ml Turbo Nuclease (Nacalai, cat. # NU0103M) in Hank’s balanced salt solution) for 7 minutes at 35 °C with gentle horizontal shaking (60 rpm). **2-** Disaggregated seminiferous tubules were left to rest in vertical position at room temperature for 2 minutes to allow tubules to decant. **3-** Supernatant was discarded and 2 ml of prewarmed (35 °C) collagenase-trypsin solution (0.55 mg/ml collagenase, 0.37 mg/ml Trypsin (Sigma, cat. # T0303), and 60 U/ml TurboNuclease in Hank’s balanced salt solution) was added. **4-** Tubules were broken down by gently pipetting 3-5 times with a transfer pipette and then incubated for 20 minutes at 35 °C with gentle horizontal shaking (60 rpm). Then, tubules were further processed by gently pipetting 3-5 times with a transfer pipette. **5-** 800 μl of serum and 50 μl of protease inhibitor (Pierce, cat. # A32965, 1 tablet in 1 ml of water) were added, mixed by inversion, and incubated on ice for 2 minutes to allow remaining tubules to decant. Cells from the supernatant were passed through a 40 μm cell strainer. At this point, a sample of 50 μl was taken to analyze spermatocyte stage by immunostaining of chromosome spread (γH2AX, SYCP1 and SYCP3 were used as markers). **6_F-_** Volume was completed to 8 ml with room temperature 1 X PBS and then 500 μl of 16 % PFA solution (Electron Microscopy Sciences, cat. # 15710) was added. Cells were incubated at room temperature with rotation for 5 minutes. **6_NF-_** For purification of spermatocytes without PFA fixation, PFA was omitted, and we proceeded straight on to Percoll centrifugation (step 11NF). **7_F-_** 1 ml of 1.25 M glycine solution was added to quench PFA, and cells were incubated for 4 min at room temperature with rotation. **8_F-_** Cell were pelleted by centrifugation at 800 g for 10 minutes 4 °C on a swinging bucket rotor. **9_F-_** The supernatant was discarded and the pellet containing cells was resuspended with 1 ml of ice-cold 1X PBS by gently pipetting until no cell clumps could be detected. **10_F-_** Ice cold 1X PBS was added to complete a total volume of 8 ml. Then, 80 μl of 1.25 M glycine was added. **11_F-_** A trimmed plastic 1 ml pipette tip was used to gently place the cell suspension on top of 4 ml of isotonic 28 % v/v Percoll solution (28 ml Percoll, plus 10 ml 10 X PBS solution a completed to a total volume of 100 ml using milliQ water) in a 15 ml tube on ice. **11_NF-_** A trimmed 1 ml plastic pipette tip was used to place the cell suspension on top of a 28:36 % v/v discontinuous isotonic-Percoll gradient (3 ml each phase) in a conical 15 ml tube on ice. **12-** The tube was centrifuged at 4 °C at 800 g for 20 minutes on a swinging bucket rotor with the brake set on “slow” or “no-brake” so that the interphase was not disrupted by the braking process. **13-** Spermatocytes were collected from PBS:28 % Percoll interphase **(F)** (Fig. 1B) or, 28:36 % Percoll interphase **(NF)** and placed on a new 15 ml tube. No more than 3 ml were collected. **14-** Ice cold 1 X PBS was added to the collected spermatocytes to complete a total volume of 14 ml and mixed by inversion. A 600 μl sample was used to assess cell purity by immunostaining of chromosome spread (γH2AX, SYCP1 and SYCP3 were used as markers, Fig. 1C). **15-** Cells were centrifuged at 800 g at 4 °C for 10 minutes and the supernatant was discarded.

### ChIP-seq

Percoll-enriched spermatocytes (93 and 80% late leptotene/early zygotene spermatocytes for replicate 1 and 2, respectively) were resuspended in 1 mL of ChIP buffer containing 5 μl of protease inhibitor cocktail (CalbioChem, cat # 539134-1) (50 mM Tris-HCl pH 8, 1 mM EDTA, 150 mM NaCl, 1% Triton X-100, 0.1 % sodium deoxycholate, 0.1 % SDS) and sonicated for 24 minutes using the Covaris E220 evolution (peak power 140, duty factor 5, 200 cycles per burst). The sample was cleared at 12,000 g for 10 min at 4 °C, and the supernatant containing chromatin was incubated with 20 μg of antibody (INO80 antibodies, abcam cat # ab105451, lot GR3206182-2) overnight at 4°C. Antibody/chromatin complexes were captured with ChIP-grade protein A/G magnetic beads (Thermo Fisher, cat # 26162) for 2 h at 4 °C, washed 2 times with increasing salt concentrations (20 mM Tris-HCl pH 8, 2 mM EDTA, 150-500 mM NaCl, 1 % Triton X-100, 0.1 % sodium deoxycholate), and once with lithium buffer (10 mM Tris-HCl pH 8, 1 mM EDTA, 250 mM LiCl , 1 % Igepal, 0.7 % sodium deoxycholate). The beads were washed twice with TE buffer pH 7.4 (10 mM Tris-HCl, 1 mM EDTA) and DNA was eluted twice by incubation at 65 °C for 30 minutes using 150 μl elution solution (1 % SDS, 100 mM NaHCO3 pH 9 made freshly). PFA cross-link was reversed overnight by adding 12 μl of 5 M NaCl and incubating at 65 °C for at least 18 hs. Next day, 6 μl of 0.5 M EDTA, 12 μl of 1 M Tris-HCl (pH 6.5) and 1 μl RNAse A (Qiagen cat # 19101) were added to the sample and incubated for 30 min at 37 °C. Then, 5 μl of proteinase K (approximately 3 U, Qiagen, cat. # 19131) were added and incubated for 1 h at 56°C. The DNA was purified using the minElute PCR purification kit (Qiagen, cat. # 28004) using 7 volumes of PB buffer (provided with the kit) and eluting twice in 12 μl of elution buffer each time. Eluted DNA was quantified using Qubit (Life Technologies) before library preparation. INO80 antibody specificity was confirmed by western blot analysis of INO80 knockout and wild type control MEFs (8). For input samples, zygotene spermatocytes were collected from synchronized mice at 8 days after retinoic acid injection. Cells were sonicated and DNA was processed as in ChIP-seq samples, except for the fact that antibodies were omitted, and no protein A/G magnetic beads purification was performed.

### Library preparation and sequencing

Library preparation and sequencing methods are detailed in Table S1. Before sequencing, samples were quantified by qPCR using Kapa library quantification kit (cat # KK4854), and size and quality of DNA was assessed using Agilent Tape station.

#### In-house library preparation method

For INO80 replicate 1 and both input samples 20 μl of the eluted DNA was incubated with 30 μl of end-repair mix (0.66 mM dNTP mix (NEB, cat. # N0447S), 100 U/ml T4 DNA polymerase (NEB, cat. # M0203L), 33 U/ml Klenow fragment (NEB, cat. # M0210S), 333 U/ml T4 PNK (cat. # M0201L), and 1.67 x T4 PNK buffer) at 20 °C for 30 minutes. DNA was purified using Qiagen Minelute kit using 7 volumes of PB buffer and eluted in 12 μl of EB buffer. 10 μl of eluted DNA were mixed with 40 μl of A-tailing mix (0.25 mM dATP, 125 U/ml Klenow fragment (cat. # NEB M02105S), and Klenow buffer 1.25 X) and incubated at 37 °C for 30 minutes. DNA was purified again using Qiagen Minelute kit using 7 volumes of PB buffer and eluted in 12 μl of EB buffer. Truseq single index adaptors (Illumina, cat. # 20015960 or 20015961) were diluted according to the DNA concentration (insert:adaptor molar ratio of 1:2, with maximal dilution of 1:50) and 1 μl of a specific diluted adaptor was added to each sample. 20 μl of ligation mix (30 U/μl T4 DNA ligase (cat. # NEB M0202L), and 1.58 X ligase buffer) was added to 10 μl of the insert:adaptor mix and incubated for 30 minutes at 20°C. DNA was purified using Qiagen Minelute kit using 7 volumes of PB buffer and eluted in 12 μl of EB buffer. Finally, DNA was amplified using the Kapa HiFi Hot Start library amplification kit (cat. # KK2621) according to manufacturer’s instructions.

#### Adaptase library preparation method

For INO80 replicate 2, ACCEL-NGS® 1S PLUS DNA LIBRARY KIT (cat. # 10024) was used in conjunction with Swift unique dual indexing kit (cat. # X9096), following manufacturer’s instructions.

### ChIP-seq data processing and analysis

#### Alignment and quality filtering

Because INO80 ChIP-seq replicate 2 was prepared for sequencing using adaptase technology, before alignment, at the beginning of each read 10 bases were trimmed for both R1 and R2 using fastp (9), as recommended by the library preparation kit manufacturer. For further steps, replicates and inputs were processed in parallel. Reads were adapter-trimmed and quality-pruned using fastp with default settings. Then, reads were aligned to GRCm38/mm10 using bwa mem (10) (Version 0.7.15) with default settings except for option ‘-M’ for Picard compatibility. Picard (version 2.21.2, http://broadinstitute.github.io/picard/) and SAMtools (version 1.11) (11) were used to obtain mapping quality metrics, remove duplicates and filter reads. Only primary alignment reads that were not duplicated, properly paired, with a MAPQ > 30, and not placed in mitochondrial chromosome were kept. MultiQC (version 1.7) was used to generate metrics reports (12). Table S2 contains INO80 ChIP-seq and input mapping statistics.

#### ChIP-seq quality control metrics

Cross-correlation analysis was done using phantompeakqualtools (version 1.2, (13)) with default parameters. Results can be found at Fig. S1. To calculate fraction of reads in peaks (FRIP) we used samtools view. The list of peaks used for each replicate was not black-greylist filtered. Mapping statistics were obtained using samtools stats (number of mapped reads) or samtools flagstat (number of uniquely mapped reads).

#### Greylist regions

Greylist regions were prepared for each input using the GreyListChIP R-package (R package version 1.24.0, https://bioconductor.org/packages/release/bioc/html/GreyListChIP.html) together with the Bsgenome.Mmusculus.UCSC.mm10 package (https://bioconductor.org/packages/release/data/annotation/html/BSgenome.Mmusculus.UCSC.mm10.html) using default settings (greyListBS command). Both lists were merged to create a single greylist, which was later used to filter peaks.

#### Reproducibility analysis

To analyze the reproducibility of replicates we analyzed how many peaks were present in both replicates. The peaks used for this analysis were called with MACS2 (version 2.2.7.1, (14)) using broad mode and default parameters. Peaks for INO80 ChIP-seq replicate 1 were called using input 1 as control and peaks for INO80 ChIP-seq replicate 2 were called using input 2 as control. Both lists of peaks were filtered for black-greylist regions before intersecting them. We also analyzed reproducibility using Irreproducible Discovery Rate (IDR) (15). To do so, we created two self-pseudo replicates of each INO80 ChIP-seq real replicate and each input by shuffling the reads and then splitting them in two equal parts. Then, self-pseudo replicate 1 of real replicate 1 was merged with self-pseudo replicate 1 of real replicate 2 to create a pooled-pseudo replicate. Same was done for replicate 2 and for both inputs. Peaks were called for all the bam files (real replicates, self-pseudo replicates and pooled replicates) using MACS2, broad mode and a lenient setting of −p 0.001. IDR analysis was done using IDR software (version 2.0.3, https://www.encodeproject.org/software/idr/) on the lists of peaks already black-greylist filtered and p-value sorted. The peaks with an adjusted IDR value >= 540 (IDR =< 0.05) were used for further analysis.

#### Peak annotation, genome and gene ontology analysis

Peaks with and IDR =< 0.05 (true replicates IDR analysis) were annotated using HOMER’s annotatePeaks.pl script ((16), version 4.10). Annotated peaks can be found at table S3. Genome ontology analysis was also done using HOMER with same annotatePeaks.pl script plus the -genomeOntology option. The results can be found at table S4. Functional annotation analysis was done using DAVID with default parameters (17,18). To generate the list of genes fed to DAVID, INO80 annotated peaks were first filtered for their distance to the closest promoter, keeping only genes with an INO80 peak closer than 5,000 bp. Further processing and plotting were done using R (https://www.r-project.org/, version 4.1.1) and packages within tidyverse (https://joss.theoj.org/papers/10.21105/joss.01686). Functional analysis results can be found at table S5.

#### Heatmaps

Bigwig coverage tracks were generated using bamCoverage tool from deepTools (version 3.4.3, (19)) with a bin size of 1 bp, no smoothing and RPKM normalization. Average aggregate profiles and heatmaps were plot using deepTools as well (computeMatrix and plotHeatmap).

#### Statistical analysis

Statistical tests were done using R (version 4.1.1) and are reported in the figure legends or main text.

#### Data availability

The datasets generated and analyzed in this study are currently available from the corresponding author. Datasets will be publicity available immediately after manuscript publication.

## Results and Discussion

We first attempted to separate unfixed spermatocytes obtained from synchronized mice testis by density gradients and centrifugation using a discontinuous percoll gradient (20) (Fig. 1B). We obtained an enriched fraction of spermatocytes (zygotene/pachytene) post Percoll gradient corresponding to 63.7 ± 11.6 % (n = 3 independent experiments) (before purification, 49.4 ± 13.1 %, n = 12). This relatively low spermatocyte enrichment obtained after Percoll purification may be caused by cell aggregation and cell lability, thus we reasoned that paraformaldehyde fixation of spermatocytes before separating by Percoll gradient may prevent aggregation and stabilize cells, resulting in increased reproducibility and improved purification efficiency. Indeed, we observed that treatment of spermatocytes (zygotene/pachytene) obtained from synchronized testis with 1% PFA resulted in improved cell enrichment (81.3 ± 11.8%, n = 55) (Fig. 1C and D). We obtained 8.7-12 ×10^6^ cells per 4-6 mice (Table 1).

**Table 1.** Analysis of zygotene cell populations after Percoll purification.

| Targeted meiotic stage | Number of mice | Number of total cells (X10<sup>6</sup>) | % targeted cell after percoll |
| --- | --- | --- | --- |
| Zygonema | 5 | 9.3 | 87 |
|  | 6 | 12 | 96 |
|  | 4 | 8.7 | 91 |

We conclude that cell treatment with PFA followed by discontinuous density Percoll gradient is an efficient and reproducible approach to enrich mouse synchronized spermatocytes at different stages of prophase I.

To test cell quality and evaluate suitability of the material for genome wide analysis we performed INO80 ChIP-seq using early zygotene spermatocytes. We chose INO80 because it is a chromatin remodeler required for normal meiotic progression (21). We reason that INO80 chromatin binding profiles may help explain the observed phenotype at the molecular level, specifically by identifying INO80 target genes or other genomic structures. Heatmaps (Fig. 2A) and visualization of specific genomic sites (Fig. 2B) obtained from two independent replicates show a robust and reproducible signal for INO80 genome wide. This was corroborated when we calculated NSC and RSC cross-correlation parameters (Fig. S1), as well as the fraction of reads in peaks (FRIP value 1 and 3 for replicate 1 and 2, respectively), which are all equal or above the minimum ENCODE-recommended standard. We detected 5,104 and 7,757 peaks in INO80 ChIP-seq replicate 1 and 2, respectively. Most peaks detected on replicate 1 were also found in replicate 2 (92%, Fig. 2C). Reproducibility was further assessed using Irreproducible Discovery Rate (IDR) analysis, as recommended by ENCODE. The number of peaks with an IDR <= 0.05 when comparing pooled-pseudo replicates was 1.63 times larger than those found when comparing true replicates. This is well under the value of 2.0, which is the maximum recommended value. We conclude that Percoll enriched spermatocyte fractions we obtained can be used to obtain robust and reproducible genome wide profiles of chromatin binding proteins.

**Figure 2.**
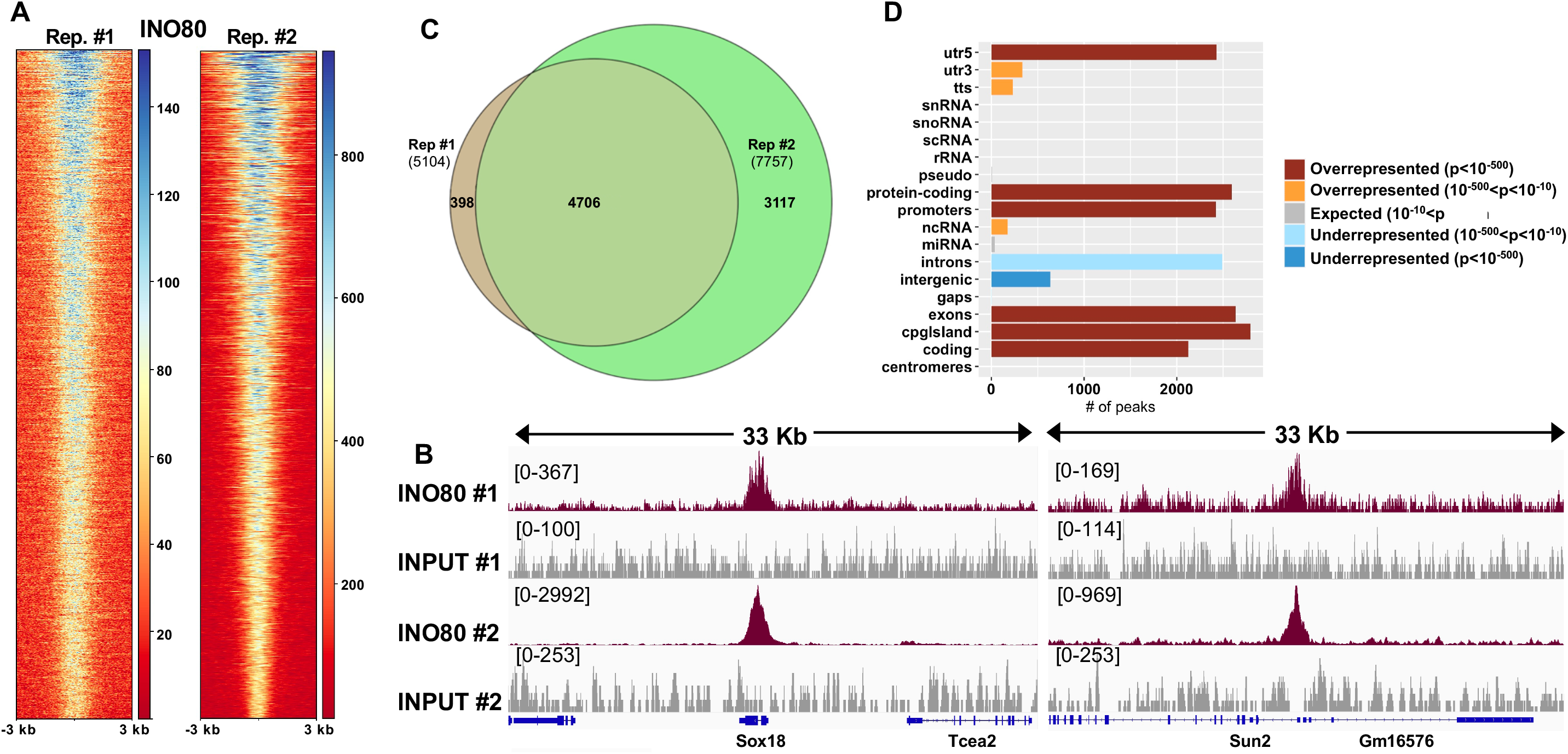
Analysis of INO80 ChIP-seq obtained from synchronized zygotene spermatocytes. **A.** Heatmap profiles corresponding to INO80 ChIP-seq signal of each replicate centered at INO80 peaks with IDR =< 0.05. **B.** Examples of INO80 ChIP-seq signals and their respective inputs at selected promoters visualized using IGV. Scales are shown between brackets at the top left corner of each sample. **C.** Ven diagram depicting the intersection of peaks of both INO80 ChIP-seq replicates. **D.** Genomic features occupied by INO80. Colors represent whether the number of peaks found at a given genomic feature is larger (overrepresented), equal (expected) or lower (underrepresented) than expected by chance.

We further analyzed INO80 chromatin binding profiles using the list of peaks that had an IDR <= 0.05 (true replicates comparison). Genome ontology analysis revealed that INO80 is found at CpG islands and protein-coding genes (promoters, exons, UTR-5’) at a much higher frequency than that expected by chance. However, INO80 has reduced presence in most non-protein-coding genes (rRNA, miRNA, snRNA, snoRNA, scRNA), intergenic regions, and centromeres (Fig. 2D and table S4). Functional analysis revealed that several genes found near an INO80 peak (less than 5,000 bp) are genes that code for proteins that regulate transcription (table S5). Indeed, “Transcription Regulation” is the second most significant Uniprot keyword associated with genes close to INO80 (389 hits and an FDR of 1.95×10^−47^), and “transcription, DNA-templated” is the third most enriched GO-Biological Process term (403 hits and an FDR of 3.95×10^−33^). Further, we searched for genes with recognized meiotic functions among genes near INO80 peaks. We used in our search a manually cured list of candidate genes as well as the genes annotated with the following GO terms: chromatin remodeling, DNA repair, meiosis I, chromosome pairing at meiosis, and synaptonemal complex assembly. We identified genes involved in critical meiotic processes such as telomere-led rapid prophase movements (SUN1 and SUN2) and homologous directed repair of double-strand break (MND1, ERCC4, and MSH2) (complete list of genes is found in table S6). Overall, our results suggest that INO80 could have a broad effect on transcription in zygotene spermatocytes, both by direct action at gene promoters as well as indirectly, by regulating transcription factors.

In summary, we have developed a methodology for enrichment of synchronized mouse spermatocytes at different stages of prophase I, that provide valuable material for experiments involving ChIP followed by massive sequencing. We predict that material obtained using this protocol will be suitable for other common genome wide approaches such as RNA-seq and ATAC-seq. We have also revealed the genome wide localization of INO80 chromatin remodeler in zygotene spermatocytes (the most advanced spermatocyte seen in INO80 knockout mouse).

## Author contributions

AC performed experiments, analyzed data, wrote, and edited the manuscript. IG performed experiments and edited the manuscript. ROC performed experiments, analyzed data, and edited the manuscript. RJP analyzed data, wrote, and edited the manuscript.

---

**Figure S1.** Cross-correlation analysis corresponding to INO80 ChIP-seq peaks.

**Table S1.** Library preparation and sequencing methods.

**Table S2.** ChIP-seq mapping statistics.

**Table S3.** Annotated INO80 ChIP-seq peaks.

**Table S4.** Genome ontology analysis.

**Table S5.** Functional annotation for INO80 ChIP-seq peaks.

**Table S6.** List of Meiosis related genes found near INO80 ChIP-seq peaks.

## References

1. Romer, K. A., de Rooij, D. G., Kojima, M. L., and Page, D. C. (2018) Isolating mitotic and meiotic germ cells from male mice by developmental synchronization, staging, and sorting. Dev Biol 443, 19–34

2. Hogarth, C. A., Evanoff, R., Mitchell, D., Kent, T., Small, C., Amory, J. K., and Griswold, M. D. (2013) Turning a spermatogenic wave into a tsunami: synchronizing murine spermatogenesis using WIN 18,446. Biol Reprod 88, 40

3. Patel, L., Kang, R., Rosenberg, S. C., Qiu, Y., Raviram, R., Chee, S., Hu, R., Ren, B., Cole, F., and Corbett, K. D. (2019) Dynamic reorganization of the genome shapes the recombination landscape in meiotic prophase. Nat Struct Mol Biol 26, 164–174

4. Chen, Y., Lyu, R., Rong, B., Zheng, Y., Lin, Z., Dai, R., Zhang, X., Xie, N., Wang, S., Tang, F., Lan, F., and Tong, M. H. (2020) Refined spatial temporal epigenomic profiling reveals intrinsic connection between PRDM9-mediated H3K4me3 and the fate of double-stranded breaks. Cell Res 30, 256–268

5. Peters, A. H., Plug, A. W., van Vugt, M. J., and de Boer, P. (1997) A drying-down technique for the spreading of mammalian meiocytes from the male and female germline. Chromosome research 5, 66–68

6. Guiraldelli, M. F., Eyster, C., Wilkerson, J. L., Dresser, M. E., and Pezza, R. J. (2013) Mouse HFM1/Mer3 is required for crossover formation and complete synapsis of homologous chromosomes during meiosis. PLoS Genet 9, e1003383

7. Guiraldelli, M. F., Felberg, A., Almeida, L. P., Parikh, A., de Castro, R. O., and Pezza, R. J. (2018) SHOC1 is a ERCC4-(HhH)2-like protein, integral to the formation of crossover recombination intermediates during mammalian meiosis. PLoS Genet 14, e1007381

8. Runge, J. S., Raab, J. R., and Magnuson, T. (2018) Identification of Two Distinct Classes of the Human INO80 Complex Genome-Wide. G3 (Bethesda) 8, 1095–1102

9. Chen, S., Zhou, Y., Chen, Y., and Gu, J. (2018) fastp: an ultra-fast all-in-one FASTQ preprocessor. Bioinformatics 34, i884–i890

10. Li, H. (2013) Aligning sequence reads, clone sequences and assembly contigs with BWA-MEM. arXiv.ORG

11. Li, H., Handsaker, B., Wysoker, A., Fennell, T., Ruan, J., Homer, N., Marth, G., Abecasis, G., Durbin, R., and Genome Project Data Processing, S. (2009) The Sequence Alignment/Map format and SAMtools. Bioinformatics 25, 2078–2079

12. Ewels, P., Magnusson, M., Lundin, S., and Kaller, M. (2016) MultiQC: summarize analysis results for multiple tools and samples in a single report. Bioinformatics 32, 3047–3048

13. Kharchenko, P. V., Tolstorukov, M. Y., and Park, P. J. (2008) Design and analysis of ChIP-seq experiments for DNA-binding proteins. Nat Biotechnol 26, 1351–1359

14. Zhang, Y., Liu, T., Meyer, C. A., Eeckhoute, J., Johnson, D. S., Bernstein, B. E., Nusbaum, C., Myers, R. M., Brown, M., Li, W., and Liu, X. S. (2008) Model-based analysis of ChIP-Seq (MACS). Genome Biol 9, R137

15. Li, Q. H., Brown, J. B., Huang, H. Y., and Bickel, P. J. (2011) Measuring Reproducibility of High-Throughput Experiments. Ann Appl Stat 5, 1752–1779

16. Heinz, S., Benner, C., Spann, N., Bertolino, E., Lin, Y. C., Laslo, P., Cheng, J. X., Murre, C., Singh, H., and Glass, C. K. (2010) Simple combinations of lineage-determining transcription factors prime cis-regulatory elements required for macrophage and B cell identities. Mol Cell 38, 576–589

17. Huang da, W., Sherman, B. T., and Lempicki, R. A. (2009) Bioinformatics enrichment tools: paths toward the comprehensive functional analysis of large gene lists. Nucleic Acids Res 37, 1–13

18. Huang da, W., Sherman, B. T., and Lempicki, R. A. (2009) Systematic and integrative analysis of large gene lists using DAVID bioinformatics resources. Nat Protoc 4, 44–57

19. Ramirez, F., Ryan, D. P., Gruning, B., Bhardwaj, V., Kilpert, F., Richter, A. S., Heyne, S., Dundar, F., and Manke, T. (2016) deepTools2: a next generation web server for deepsequencing data analysis. Nucleic Acids Res 44, W160–165

20. Kogo, H., Kowa-Sugiyama, H., Yamada, K., Bolor, H., Tsutsumi, M., Ohye, T., Inagaki, H., Taniguchi, M., Toda, T., and Kurahashi, H. (2010) Screening of genes involved in chromosome segregation during meiosis I: toward the identification of genes responsible for infertility in humans. J Hum Genet 55, 293–299

21. Serber, D. W., Runge, J. S., Menon, D. U., and Magnuson, T. (2016) The Mouse INO80 Chromatin-Remodeling Complex Is an Essential Meiotic Factor for Spermatogenesis. Biol Reprod 94, 8

