## Supplementary figures and images for "Efficient enrichment of synchronized mouse spermatocytes suitable for genome-wide analysis"

### Supplementary Figure 1

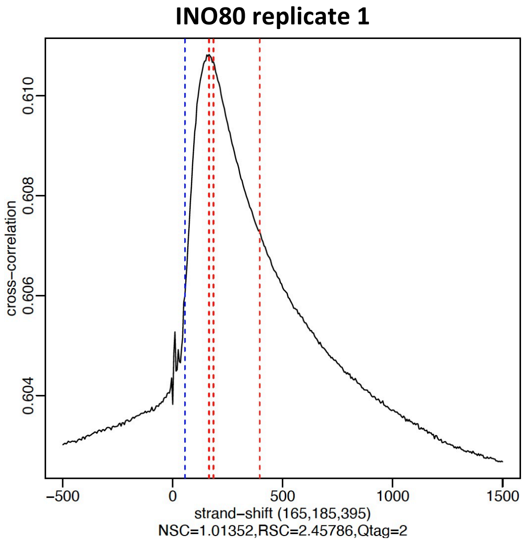

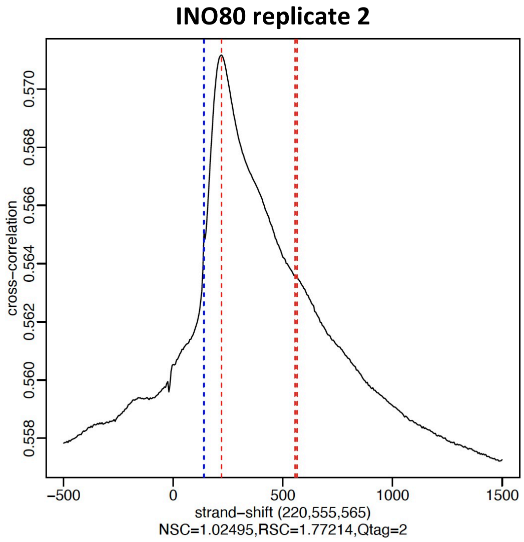
